# Using connectivity to identify climatic drivers of local adaptation

**DOI:** 10.1101/145169

**Authors:** Stewart L. Macdonald, John Llewelyn, Ben L. Phillips

## Abstract

*This preprint has been reviewed and recommended by Peer Community In Evolutionary Biology (<u>http://dx.doi.org/10.24072/pci.evolbiol.100034</u>).* Despite being able to conclusively demonstrate local adaptation, we are still often unable to objectively determine the climatic drivers of local adaptation. Given the rapid rate of global change, understanding the climatic drivers of local adaptation is vital. Not only will this tell us which climate axes matter most to population fitness, but such knowledge is critical to inform management strategies such as translocation and targeted gene flow. While simple assessments of geographic trait variation are useful, geographic variation (and its associations with environment) may represent plastic, rather than evolved, differences. Additionally, the vast number of trait–environment combinations makes it difficult to determine which aspects of the environment populations adapt to. Here we argue that by incorporating a measure of landscape connectivity as a proxy for gene flow, we can differentiate between trait–environment relationships underpinned by genetic differences versus those that reflect phenotypic plasticity. By doing so, we can rapidly shorten the list of trait–environment combinations that may be of adaptive significance. We demonstrate how this reasoning can be applied using data on geographic trait variation in a lizard species from Australia's Wet Tropics rainforest. Our analysis reveals an overwhelming signal of local adaptation for the traits and environmental variables we investigated. Our analysis also allows us to rank environmental variables by the degree to which they appear to be driving local adaptation. Although encouraging, methodological issues remain: we point to these issue in the hope that the community can rapidly hone the methods we sketch here. The promise is a rapid and general approach to identifying the environmental drivers of local adaptation.

## Introduction

It is only recently that we have begun to appreciate the speed with which evolution can happen; not only over relatively short timespans (e.g., 1, 2–4), but also at small spatial scales (5). Rapid local adaptation has been recorded in response to a wide suite of environmental drivers, including invasive species, and pollution (6). We expect climate to also be a major driver of local adaptation (e.g., 7, 8), and understanding the way in which species respond to climate is of increasing importance because anthropogenic climate change is proceeding at such a rate that there are concerns that many species will be unable to evolve rapidly enough to avoid extinction (9, 10).

Evolution typically optimizes phenotypes, but the optimum will vary through both time and space (11, 12), in turn leading to populations ('demes') that have, on average, higher fitness in their home environment than an immigrant would: local adaptation. While adaptive optima for traits almost always vary geographically, it does not follow that all geographic trait variation is due to local adaptation. Geographic trait variation can arise due to other factors, such as phenotypic plasticity (including developmental plasticity and maternal effects), neutral clines, and environmental factors (such as geographic variation in fitness-reducing parasites). These factors can give the appearance of local adaptation (10, 11), complicating our identification of climate-relevant adaptive variation.

To circumvent these issues, evolutionary biologists use experimental approaches to demonstrate local adaptation (12, 13). Experiments designed to detect local adaptation typically utilise one of two techniques: 1) reciprocal transplants, which are done *in situ*, and are considered the gold standard for demonstrating local adaptation; or 2) common garden experiments, which are usually done in the lab where it is easier to control each environmental variable (12). Both of these techniques can be difficult, for reasons of time, expense, logistics, or ethics. This difficulty increases as the number of separate demes and environmental variables to be tested increases and as the generation time of the organism increases (12). Additionally, although reciprocal transplants will detect signs of local adaptation, they are not necessarily suited to identifying the environmental drivers of that local adaptation (14). This is because *in situ* reciprocal transplants necessarily encompass all the environmental variables that differ between the transplant locations. Lab-based common garden approaches may, in principle, be more suited to identifying environmental drivers (because the environment may be under a degree of control), but in practice it often remains impossible to identify the environmental drivers of trait variation seen in the wild. Thus, the best experimental tools we have for studying local adaptation are demanding in terms of time and cost, and are unsuitable for assigning environmental drivers (such as climate variables) to adaptive variation. If we are looking for climate-driven local adaptation, this is a problem: we want to know which climate variable or variables are the main drivers of adaptation, and we urgently need this information for many species.

By definition, local adaptation has a genetic basis and is consequently weakened by high levels of gene flow (11, 15, 16). Demes with excessive inward gene flow are therefore likely to be less optimally adapted, causing an observable mismatch between optimal and actual phenotypes. Some examples of this are birds dispersing and producing clutch sizes that are not optimised for the habitat quality in which they are now nesting (17), larval salamander colouration not matching streambed colouration due to high levels of gene flow from nearby but predator-free streams (18), and stick insects in smaller habitat patches having non-cryptic colouration when the surrounding patches are larger and environmentally dissimilar (19). These observations of “migrant load” suggest an alternative technique for identifying and assessing local adaptation. First, we look across populations for relationships between the environment (e.g., mean annual temperature) and traits (e.g., morphology, physiology). By themselves, these relationships are not sufficient evidence of local adaptation — they could also be caused by phenotypic plasticity. Second, knowing that local adaptation is hindered by gene flow, we can look at whether gene flow diminishes the environmental effect. With some caveats (discussed below), in cases where data on gene flow are absent (which is often the case), habitat connectivity can be used as a substitute for gene flow. Trait–environment relationships that are strong, but which are also weakened by connectivity, are indicative of trait–environment relationships that have a genetic basis. In a statistical model, this idea would be represented as follows:

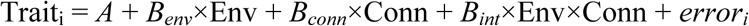

Where:

trait_i_ = trait value for individual *i*

*A* = intercept

*B*_*env*_ = coefficient of the environmental variable

Env = environmental variable (e.g., annual mean temperature) at the individual’s site

*B*_*conn*_ = coefficient of connectivity

Conn = connectivity at the individual’s site

*B*_*int*_ = coefficient of the interaction between Env and Conn

*error*_*i*_ = deviation between expected value and trait value of individual *i* Which, with slight rearrangement, can also be expressed as:

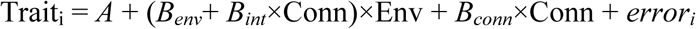

showing that the slope of the relationship between the trait and the environment now depends on the connectivity value. When the signs of B_env_ and B_int_ are in opposition, then we have a situation in which the relationship between the trait and the environment diminishes as connectivity increases.

If we now collect data on a large number of trait–environment relationships, and their interaction with connectivity, we can imagine several possible patterns emerging. These possibilities are depicted in Figure 1. Each panel represents a possible relationship between trait–environment coefficients (along the x-axis) and the interaction between environment and connectivity (y-axis). Panel A shows a set of trait–environment relationships that vary in strength, but that are not influenced by connectivity (i.e., no environment–connectivity interaction). This pattern is indicative of a system in which trait–environment relationships are predominantly driven by plastic responses of traits to their environment (i.e., traits always match the local environment, regardless of the level of inward gene flow). Panel B shows a system in which trait–environment relationships are eroded by connectivity: increased connectivity diminishes the relationship between the environment and the trait. In this situation, the interaction between the environmental variable and connectivity is negative when the environmental coefficient is positive (i.e., greater connectivity causes the environmental coefficient to decrease towards zero; bottom-right quadrant), and positive when the environmental coefficient is negative (i.e., greater connectivity causes the environmental coefficient to increase towards zero; top-left quadrant). This is the pattern we would expect if there is a genetic basis to the trait– environment relationship, such as is exhibited by local adaptation. Panel C shows the situation where the effect of the environment tends to be enhanced by connectivity. This pattern might arise in organisms that are highly mobile and can actively move to their ideal environment, thus avoiding the selective pressures that would lead to local adaptation.

**Figure 1.**
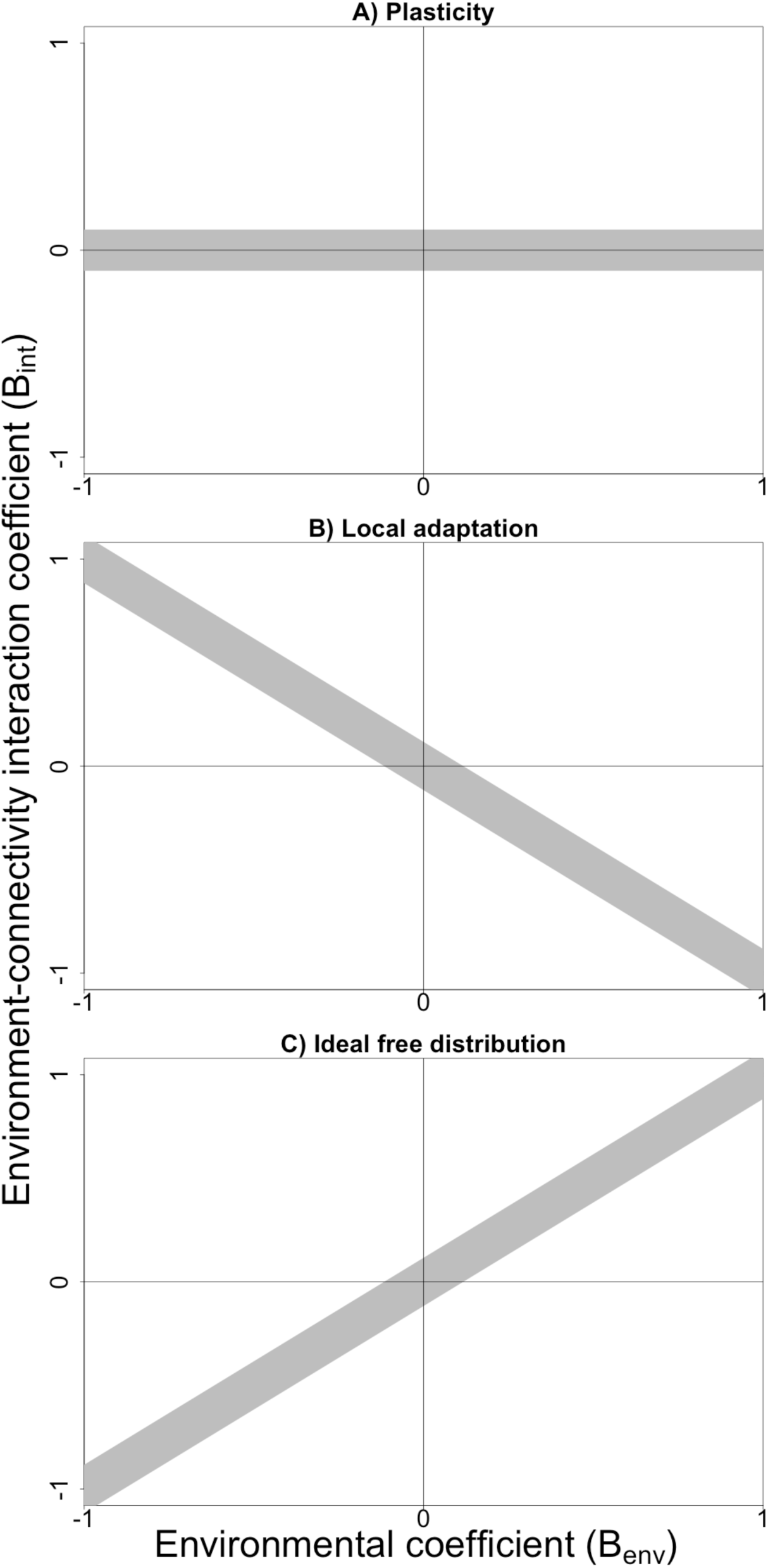
Graphs showing the concepts illustrated by plotting a set of trait–environment coefficients (e.g., the coefficient from a linear model examining the effect of annual mean site temperature on the sprint performance of organisms from that site: *B*_*env*_, x-axis) and the corresponding environment–connectivity interaction coefficients (*B_int_,* y-axis). Broad grey line represents the approximate area in which these points would fall. A) Phenotypic plasticity is suggested when trait–environment relationships are strong, but are not influenced by connectivity. B) Local adaptation is suggested when increasing connectivity diminishes the relationship between the environment and the trait. C) The effect of the environment is enhanced by connectivity. This latter pattern might arise in organisms that are highly mobile and can actively move to their ideal environment, thus avoiding the selective pressures that would lead to local adaptation.

Understanding how species respond to specific aspects of their environment is vital if we are to have any hope of halting the current rapid loss of biodiversity. Climate change is undoubtedly one of the biggest threats to global biodiversity (20, 21), and conservation biologists are looking to a variety of techniques to assess and help mitigate the impacts of climate change on vulnerable species (22–24). One technique that is likely to see increasing use is targeted, or assisted, gene flow [TGF; for review, see (22, 25)]. This technique involves the spatial redistribution of long-standing adaptations, and acts to increase genetic diversity in recipient populations, thereby bolstering capacity for evolutionary adaptation (10, 22, 24, 25). When applying TGF to help species adapt to climate change, we need to find an existing location that matches the future climate at our recipient site, and then translocate animals from that source location. It is a simple idea, but climate is multidimensional and species will not be adapting equally to each climate axis: is a difference of 0.5°C in mean temperature more important than a difference of 100mm in annual rainfall? The answer depends upon which aspects of climate (hereafter “climatic axes”) have the strongest influence on fitness.

Here we explore the idea of using connectivity to infer local adaptation. To do this we develop a case study of a lizard species from northern Australia. We use this system to examine the relationship, across sites, between traits and climatic variables. We assess how habitat connectivity affects these relationships and use the interaction between the environmental variable and connectivity to rank trait–environment combinations. In doing so, we reveal a set of trait–environment relationships that appears to be dominated by local adaptation.

## Methods

### Study species and site selection

The Rainforest Sunskink (*Lampropholis coggeri*) is a small (snout–vent length up to 45 mm), diurnal scincid lizard restricted to the rainforests of the Wet Tropics region of northeastern Australia (26). The rainforests of this region cover a wide range of environmental conditions, spanning significant elevation (0–1600 m ASL), precipitation (annual mean precipitation of 1432–8934 mm, not including input from cloud stripping), and temperature (annual mean temperature of 16.3–25.8°C) gradients. This heliothermic skink is active year-round, often seen basking in patches of sunlight on the rainforest floor. Lizards were captured by hand from sites that were selected to maximize the environmental heterogeneity sampled (Fig. 2).

**Figure 2.**
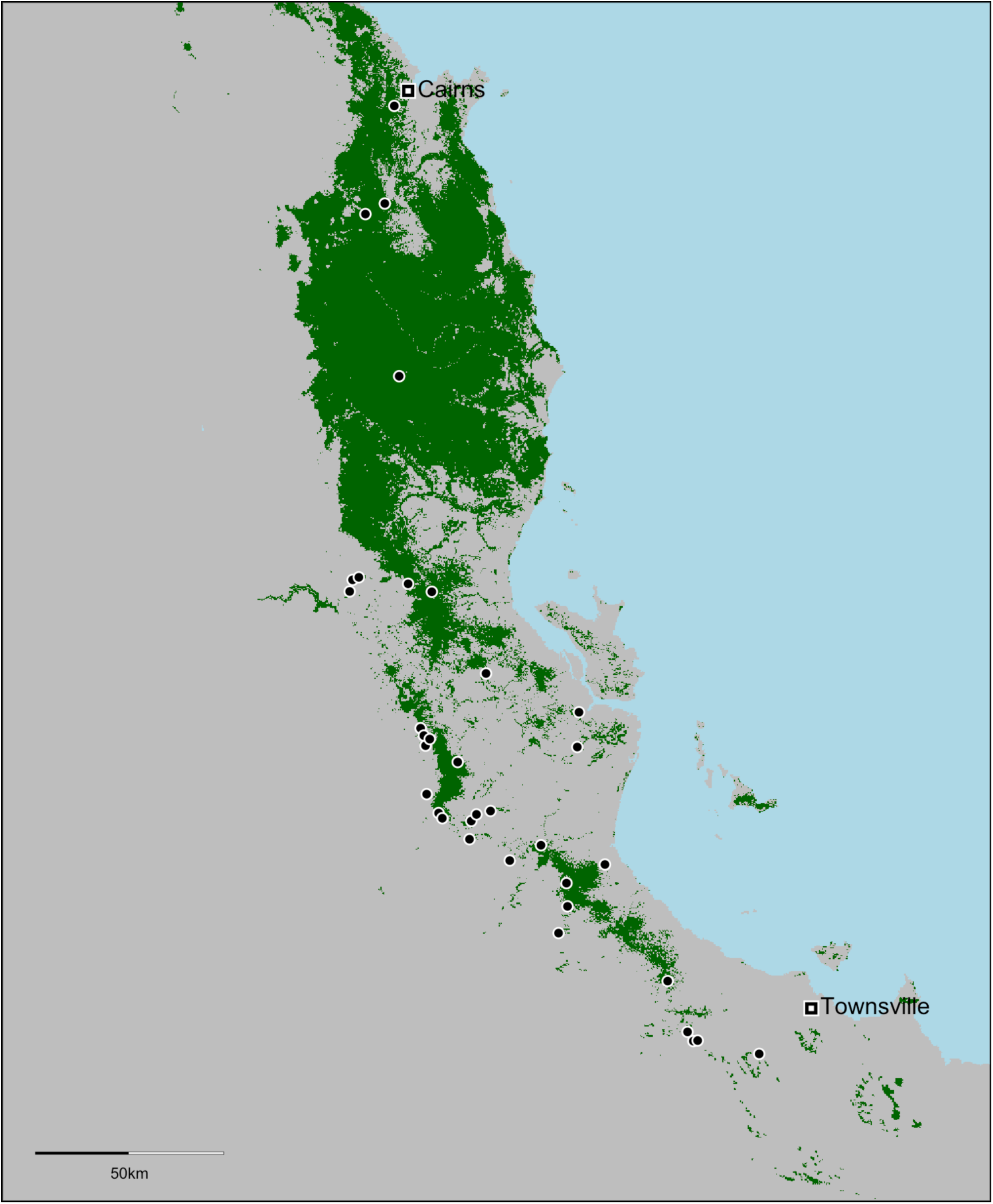
Map of the southern Australian Wet Tropics bioregion, showing the distribution of rainforest in green and the sampling locations as black dots.

Morphological measurements were obtained from 532 skinks from 32 sites. Physiological measurements were obtained from a smaller subset of these lizards: 259 skinks from 12 sites. At each site, 8–20 skinks were caught per collecting trip. Following capture, skinks were transported to James Cook University (JCU) in Townsville for trait measurement. All procedures involving lizards were approved by the JCU animal ethics committee (projects A1976 and A1726).

### Physiological trials

Physiological trials commenced within seven days of skinks being collected from the field; skinks being used only for morphology were measured and released back at their point of capture within seven days. The following measures were taken from each skink (n = 259) during laboratory trials: critical thermal minimum (CTmin), critical thermal maximum (CTmax), thermal-performance breadth for sprinting (breadth80), maximum sprint speed (Rmax), temperature at which sprint speed is optimized (Topt), active body temperature as measured in a thermal gradient (Tactive), and desiccation rate (des) (see Table S1 for further details). Details of trait measurement procedures are detailed elsewhere (see 27, 28).

### Morphological measurements

The following measurements were taken from each skink (n = 532) using digital calipers: head width (HeadW); head length (HeadL); interlimb length (ILimbL); hindlimb length (HindLL). Left and right measurements were averaged to obtain one measurement for that trait. We also recorded snout–vent length (SVL), total length, and mass (see Table S1 for further details). All measurements were taken by one person (SLM) to minimize observer bias. All morphological variables were log-transformed prior to regression analyses.

### Climatic variables, and connectivity

Because our study aimed to assess adaptation to local climate, various temperature and precipitation variables were extracted for each site (see Table S2 for details). We considered both means and extremes. It is important to consider climatic extremes, because temperature extremes may be increasing faster than mean temperatures (29), and selection may often occur during extreme weather events (30). Many environmental variables are highly correlated (27), so only the less-derived variables were used in analyses, specifically: annual mean precipitation (AMP); seasonality of precipitation (Pcov); precipitation of the driest quarter (Pdry); annual mean temperature (AMT); coefficient of variation of temperature (Tcov); average minimum daily temperature (Tmin); average maximum daily temperature (Tmax); average variance of daily maximum temperature (TmaxVar); and average variance of daily Tmin (TminVar).

Our connectivity index was designed to capture the flux of individuals through a location and is detailed in (31). Briefly, it is a measure of habitat suitability for our focal skink species, averaged over space using a species-specific estimate of dispersal potential. This approach is reasonable for any species exhibiting diffusive dispersal, and similar techniques (though different spatial-weighting functions) can be used for species exhibiting non-diffusive dispersal. As our species is an obligate rainforest-dweller, grid cells in the landscape that are rainforest and that are surrounded by rainforest have high connectivity indices, while grid cells of rainforest surrounded by non-rainforest matrix have low indices. See Table S2 for further details on all variables, and Figure S1 for correlations between all variables.

### Analysis

Our analysis aimed to assess: 1) the relationship, across sites, between each trait and each environmental variable; and 2) how connectivity affected each of these relationships (i.e., the interaction between connectivity and environment). To allow comparison of coefficients across variables, and to make interaction effect-sizes meaningful, all trait and environmental variables were standardized so they had a mean of 0 and a standard deviation of 1. Linear models were fitted for each pair of environment–trait variables, with all models including the effect of lizard body size and sex, as well as the interaction between environment and connectivity:

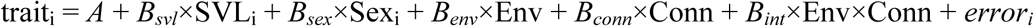

Where:

trait_i_ = standardized trait value of interest for lizard *i*

*A* = intercept

*B*_*svl*_ = coefficient of SVL SVL = lizard snout–vent length, to control for effect of body size

*B*_*sex*_ = coefficient of Sex

Sex = lizard sex (this species is sexually dimorphic in some morphological traits)

*B*_*env*_ = coefficient of environmental variable

Env = environmental variable (e.g., annual mean temperature) at the lizard's site

*B*_*conn*_ = coefficient of connectivity

Conn = connectivity index at the lizard's site

*B*_*int*_ = coefficient of interaction between Env and Conn

*error*_*i*_ = deviation between expected value and trait value of lizard *i*

A score for ranking the strength of local adaptation (*L*) was then calculated as:

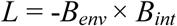

If the signs of the two coefficients (*B*_*env*_ and *B*_*int*_) are opposite (which indicates an trait–environmental relationship that is diminished by increasing connectivity, i.e., evidence for local adaptation), *L* will be positive. If the signs are the same (which indicates an environmental effect being enhanced by increased connectivity, a situation not consistent with local adaptation), *L* will be negative. Thus, higher numbers on this scale equate to stronger evidence for local adaptation in that environment–trait pair. This score can, in theory, range from –∞ to +∞. Once many environment–trait combinations have been assessed, the coefficients for all pairs can be plotted (see Fig. 1). As described in the Introduction, in a system dominated by local adaptation, we expect to see a negative relationship between *B*_*env*_ and *B*_*int*_ (Fig.1B). All analyses were conducted in R v3.2 (32).

## Results

There was substantial variation in the effect of environment (*B*_*env*_) and its interaction with connectivity (*B*_*int*_) across climate and trait variables, with *B*_*env*_ ranging from −1.8 to 1.61, and *B*_*int*_ ranging from −0.73 to 0.78 (Fig. 3). Despite this variation, a clear pattern is evident, with most points in Figure 3 appearing in the top-left or bottom-right quadrants: the quadrants in which the two coefficients have opposing signs, and where we would expect points to fall if trait–environment relationships have a genetic basis. Across these trait–environment combinations there is a distinct negative linear trend (slope= −0.36, p < 0.001). It is especially noteworthy that the trait–environment pairs with the largest coefficients are in the two quadrants indicative of local adaptation.

**Figure 3.**
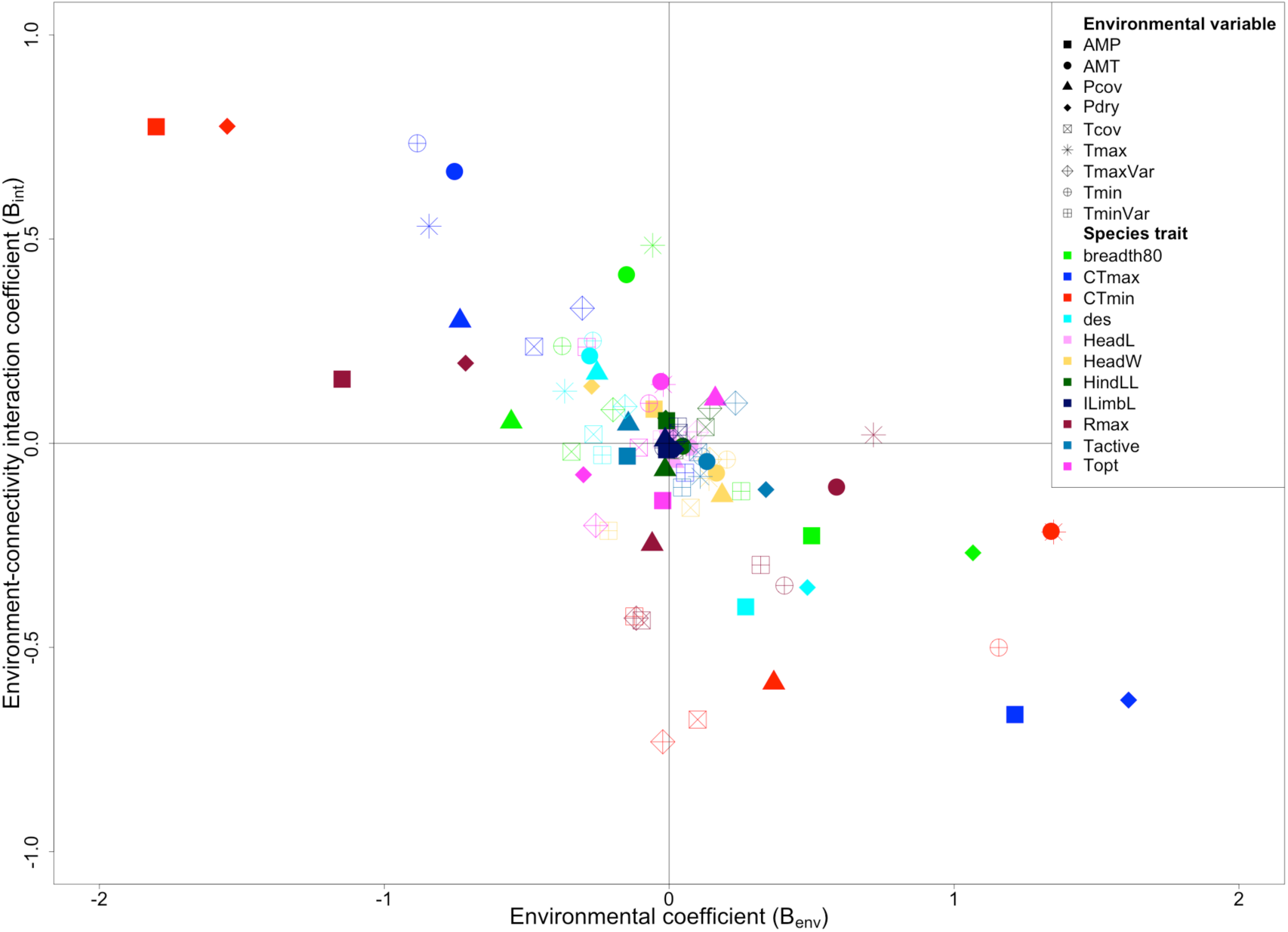
Scatterplot showing the results of 99 linear models run to assess the relationship between each trait–environment pair, and the environment–connectivity interaction. Trait– environment coefficients (*B*_*env*_) are on the x-axis, and environment–connectivity interaction coefficients (*B*_*int*_) are on the y-axis. Local adaptation is suggested when these two parameters are opposite in sign: in trait–environment pairs in which a strong environmental effect is eroded by increasing connectivity.

Overall, physiological traits showed substantially stronger environmental effects (i.e., larger values of *B*_*env*_) than did morphological traits, with the largest environmental effects being exhibited by CTmin (AMP: −1.80; Tmax: 1.35; Pdry: −1.55) and CTmax (Pdry: 1.61; AMP: 1.21). Physiological traits also showed stronger interactions between environmental effects and connectivity, again with CTmin and CTmax showing the largest interactions. These trends are apparent when we examine our index of local adaptation, *L*. Figure 4 shows a heatmap of all trait–environment pairs, ranked via reciprocal averaging according to the strength of their local adaptation index. The trait–environment pairs that show the strongest signature of local adaptation appear at the top-left in red. There is a rough divide, with most of the physiological traits on the left and most of the morphological traits on the right. The exceptions are the physiological traits Topt and Rmax, which appear at the far right of the figure.

**Figure 4.**
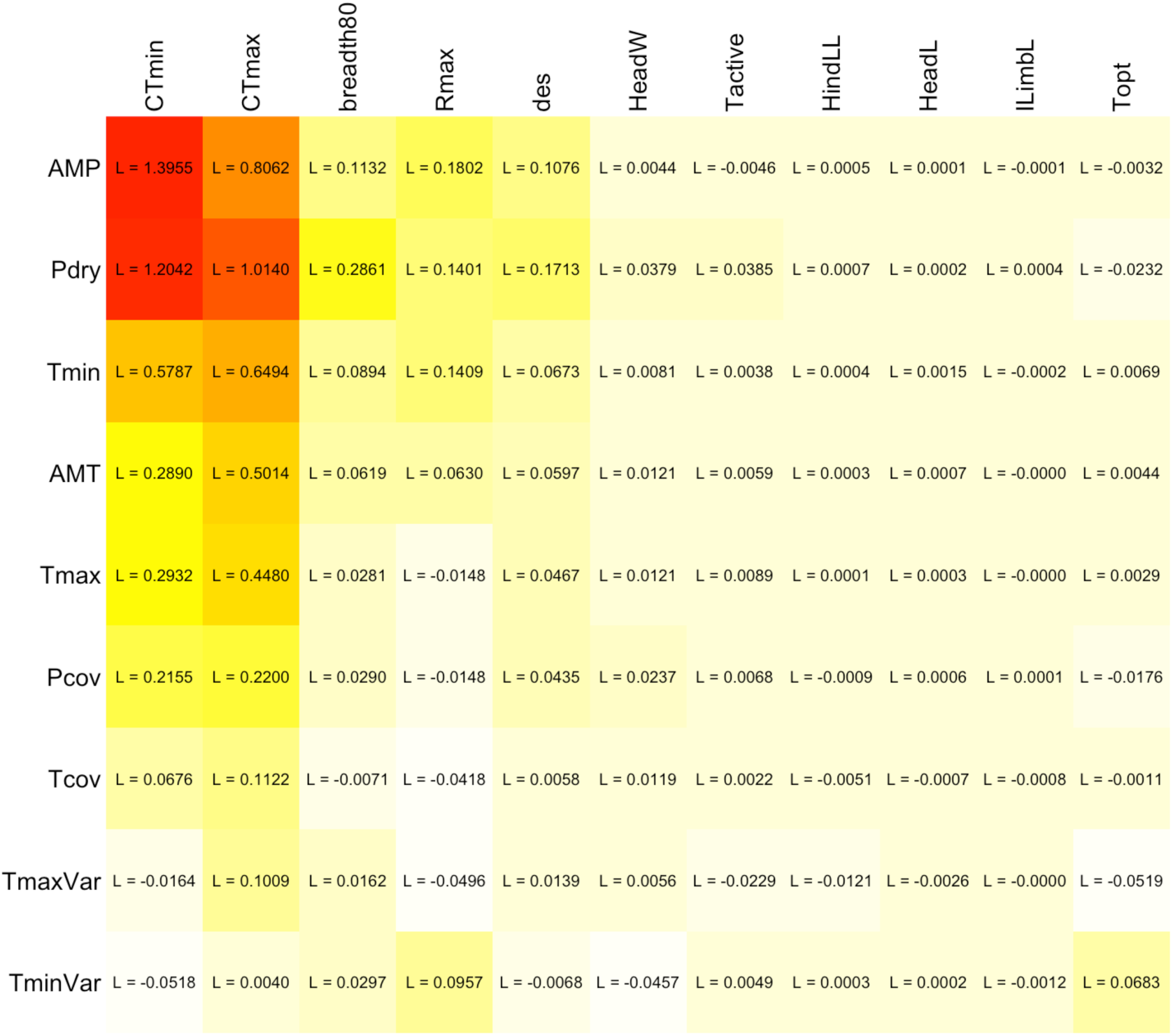
Heatmap showing the relative rankings of climate variables (rows) and morphological and physiological traits (columns). The matrix has been sorted (by reciprocal averaging) and coloured according to the strength of local adaptation, with higher values coloured red and being sorted to the top/left. See Tables S1 and S2 for explanations of the trait and environmental variables used. *L* = local adaptation index: -*B_env_*× *B_int_*

The two environmental variables that produced the strongest effects (topmost rows in Fig 4) were both precipitation related: annual mean precipitation (AMP) and precipitation of the driest quarter (Pdry). In our system, AMP and Pdry are both highly correlated with connectivity (see Fig. S1). This is expected, because our connectivity index is largely a measure of where rainforest is, and the distribution of rainforest in our study region is driven to a large degree by rainfall.

## Discussion

Understanding relationships between traits and the environment will help us plan management strategies, such as targeted gene flow (TGF), that can mitigate the impact of climate change on vulnerable species. Numerous studies have looked for (and found) trait–environment relationships (e.g., 18, 19, 33, 34–36), but the interpretation of these associations is plagued with uncertainty: are they associations due to local adaptation, neutral clines, habitat choice, or plasticity? By acknowledging that gene flow undermines adaptation, we can incorporate connectivity (a proxy for the flux of genes) into our analysis, and in doing so, separate those relationships due to fixed genetic differences, from those due to plasticity or habitat choice.

### Local adaptation

In the trait–environment combinations we assessed, physiological traits typically showed a substantially stronger effect of environment (*B*_*env*_) than did morphological traits, with the largest environmental effects shown in CTmax and CTmin (Figs. 3 & 4). Physiological traits also generally showed stronger environment–connectivity interactions (*B*_*int*_), again with CTmin and CTmax showing the largest interactions. Overall, physiological traits generally showed stronger evidence of local adaptation than did morphological traits. This result is intuitive: we would expect an ectotherm's physiological traits to be under strong selection from climate (37–39), but the fitness link between morphology and climate is much less clear. Had we also included some environmental variables that had a clearer bearing on morphology, we might have detected stronger trait–environment relationships for morphology. For example, skinks that occur in rockier habitats show various morphological adaptations to that environment (40). Including a measure of rockiness in our set of environmental variables might have allowed us to detect a signal of local adaptation for limb length. Here, however, our focus is on climatic aspects of the environment.

Of the environmental variables used, our analysis suggests that precipitation is a very strong driver of local adaptation, even in thermal traits that might not seem obviously related to precipitation (e.g., CTmin, CTmax). Although this may seem a surprising result, precipitation has been shown to directly affect growth rate, body temperature, activity patterns, and thermoregulatory opportunities in lizards (38, 41–45). Wetter areas also have higher thermal inertia (and so lower cyclical thermal fluctuations (46)), and changed environmental variance in temperature potentially has a strong influence on thermal limits (47). Additionally, Bonebrake and Mastrandrea (48) found that changes in precipitation can significantly affect modeled fitness and performance curves. Finally, comparative analyses also suggest that precipitation can influence thermal traits in many species (38). Thus, although the mechanisms linking precipitation to thermal limits are diffuse and poorly resolved, they do exist, and our analyses suggest that precipitation is a strong driver of local adaptation at thermal physiological traits.

Our analysis also suggests that temperature is an important driver of local adaptation in this system, but that extremes of temperature (encapsulated in minimum and maximum temperatures) are at least as strongly associated with local adaptation as is mean temperature. Again, this result is intuitive (natural selection from climate is likely stronger during extreme events than during normal daily temperatures) and agrees with results of empirical studies (38). Finally, the environmental variables with the weakest signals of local adaptation are Tcov (temperature seasonality), TminVar, and TmaxVar (variance of minimum and maximum daily temperatures, respectively). These variables represent predictable environmental variation occurring within an individual's lifespan and so are variables to which we might expect individuals to develop plastic responses, rather than fixed differences; local adaptation to these variables would likely be reflected in reaction norms, rather than point values for traits. (49–51).

### System-wide signal of local adaptation

The clear negative linear trend displayed in Fig. 3 is precisely what we would expect in a set of trait–environment combinations dominated by local adaptation. Migrant load (the negative effect of the immigration of less-locally adapted individuals) scales positively with immigration as well as with the strength of selection [see equation 5 in Polechová, Barton and Marion (52)]. The reason for this is that, when the strength of selection is moderately high, the environment will have a large effect on relevant traits, and therefore any immigrants coming from differing environments will be particularly maladapted and will therefore have a large and negative impact on the local phenotype. Thus, we expect trait–environment combinations with strong local adaptation to show strong effects of connectivity on the trait–environment relationship (52).

By setting up a statistical model in which the trait–environment relationship is altered by connectivity, we have allowed the possibility that the trait–environment relationship could be reversed as connectivity increases. Such an outcome is absurd from a theoretical perspective. In practice, however, our interaction coefficients were typically estimated to be around 0.36 times as strong as the main effect of environment. In this situation, reversal would only happen when connectivity values were more than 2.7 standard deviations beyond the mean (a situation that is exceedingly rare). Thus, encouragingly, our system wide analysis consistently provides parameter estimates that are theoretically sensible, despite there being no constraint within the model for them to be so.

We used long-term climatic averages and found strong evidence that local adaptation dominates over plasticity in our trait–environment set. If we had included different environmental variables, such as the conditions the lizards had recently encountered, signals of plasticity may have been more apparent. Clearly environmental variables that are similar across generations should lead to local adaptation, while environmental variables that fluctuate within generations should have a strong influence on phenotypic plasticity.

### Phenotypic plasticity

The importance of accounting for phenotypic plasticity is, however, exemplified in our dataset by the relatively strong effect of precipitation of the driest quarter (Pdry) on the temperature at which maximum sprint speed is achieved (Topt) and on maximum sprint speed (Rmax) itself. On their own, these strong trait–environment relationships might be interpreted as evidence for local adaptation. Our analysis, however, suggests that the environmental effect is largely independent of connectivity, implying that variation in these traits is due to plasticity rather than genetic differentiation. Other work (27) has shown little temporal variation in Topt (within generations) despite clear geographic variation and this, together with our results, suggests that this trait undergoes developmental plasticity, but is fixed in adult lizards. In principle, this non-effect of connectivity could also arise due to selection that is so strong that it maintains local adaptation despite high levels of gene flow [i.e., immigrants are selected against so strongly that they do not contribute to the recipient population (11)]. The trait–environment relationships for Topt and Rmax are, however, weaker than those for some other traits (e.g., CTmax and CTmin) that show clear effects of connectivity, so extremely strong selection seems an unlikely explanation for the pattern we see here.

The generally weak evidence for plasticity in our dataset should not be considered weak evidence for plasticity in these traits. Indeed many of the physiological traits we use (e.g., CTmax) are notoriously plastic, responding reversibly on timescales ranging from hours to months (53, 54). That we do not see signals of plasticity in these traits here reflects our choice of environmental variables: long-term climatic variables, rather than short-term weather variables (such as the temperature in the week before an animal was collected). We chose these long-term variables precisely because we are interested in unearthing patterns of local adaptation, rather than patterns due to rapid, reversible plasticity.

### Caveats and challenges

Our intent here has been to point out the additional inference that can be drawn from data on geographic trait variation if we account for the effect of gene flow on trait differentiation. The idea that local adaptation is eroded by gene flow offers a novel way to identify the environmental drivers of local adaptation. Such a capacity is of fundamental interest, and is also sorely needed if we are to effectively manage the impacts of climate change. The methods we use here are, however, embryonic, and in the following we point out caveats and challenges for future work.

#### Gene flow and connectivity

Our approach requires a measure of gene flow across a landscape. Here we have used environmental connectivity as a proxy for gene flow. We chose connectivity because it can be calculated relatively easily for many species by using broad scale habitat mapping datasets [e.g., vegetation mapping from DERM (55)]. Of course, these measures of connectivity should be calculated at a scale relevant to the scale of dispersal of the species in question [as ours was, using dispersal rate data for *Lampropholis coggeri* from Singhal and Moritz (56)]. While connectivity measures will often correlate with gene flow, e.g., (57)], a measure of gene flow, rather than the flow of individuals, would be preferable. Such measures are increasingly becoming available with the rise of landscape genomics tools (e.g., 58), but may still be cost-prohibitive in many cases.

While there may be better measures of gene flow, our inference might also be improved by taking into account landscape heterogeneity in the environment. Gene flow, per se, does not erode local adaptation. Rather it is an influx of maladapted genes that erodes local adaptation. Thus, a better index of this “migrant load” may well be one in which connectivity is multiplied by a measure of environmental heterogeneity, where connectivity and heterogeneity are calculated over the same spatial scale (e.g., 31). An index such as this should, in principle be a better measure of migrant load than our simple measure of connectivity. The cost, however, is that this index would need to be calculated in a standardized manner for every environmental variable under consideration.

Clearly connectivity is an imperfect measure of migrant load. By using it, we implicitly assume that all migrants are equally maladapted and have equal fitness in the recipient population. Nonetheless, connectivity should scale positively with migrant load, and our analysis using simple connectivity generated a coherent and intuitively sensible result. This is encouraging, suggesting that, in the absence of precise estimates of migrant load, a readily calculable connectivity metric may suffice to elucidate broad patterns.

#### Linear trait–environment relationships, and covariation with connectivity

Our method assumed that traits have a linear relationship to the environment (at least at the environmental scale across which we are looking). In many instances, this will be a reasonable null assumption: it seems unlikely, for example, that a trait such as desiccation resistance would be high in dry environments, low in moderately wet environments, and then high again in very wet environments. The assumption bears particular mention, however, in the situation where the connectivity index is strongly correlated with one or more of the other environmental variables being used. In our system, for example, AMP and Pdry are correlated with connectivity (Fig. S1). Where the environment–connectivity correlation is very strong, the interaction term in our model (Conn×Env) could be interpreted as a quadratic term for environment (i.e., Env^2^). In these cases, it is possible that a strong connectivity interaction is, in fact, pointing to a non-linear trait–environment relationship. Thus, for environmental variables that correlate with connectivity (and there will always be some), careful consideration needs to be given to the possibility of a quadratic fitness function between trait and environment. In our case, it remains possible, for example, that the strong influence of precipitation on local adaptation in our system is spurious, and instead reflects non-linear relationships between optimal trait values and precipitation. We can, however, think of no obvious reason why thermal limits should respond quadratically to precipitation, nor why desiccation rates and other physiological traits should also do so. Thus, in our case, we are inclined to accept the importance of this environmental variable in driving local adaptation in our system.

#### Covariation between explanatory variables

As in any multiple regression analysis, our capacity to make precise coefficient estimates diminishes if there is substantial covariation between our explanatory variables. If a sampling regime is being designed *de novo*, care should be taken to sample sites in such a way that covariation between environmental variables (including connectivity) is avoided as far as possible. Such an aim can be achieved by, for example, strategically exploiting latitudinal and altitudinal gradients.

#### Multivariate traits and environments

Here we examined one trait–environment combination at a time. Doing so may potentially miss relationships that only appear in multivariate analyses. For example, if two environmental variables are negatively correlated but both have a positive effect on a trait, it is possible that these countergradients can obscure the univariate relationship. Similar problems are encountered when examining response to selection over time (59) and, with our approach, may lead us to underestimate the number of important environmental drivers of local adaptation. Analysis incorporating multiple environmental predictors is possible, but such a model will rapidly become saturated with parameters. To minimize the problem of countergradients, again, care should be taken to sample environmental spaces in such a way as to minimize correlations between environmental variables.

An additional analytical challenge is to treat traits as multivariate. Here we have treated each measured trait as independent. In reality, however, traits covary and this covariance can have both genetic and environmental origins (60). As a corollary, selection acts on the multivariate trait, and causes populations to move in multivariate trait space (61). Consequently, local adaptation perhaps should be measured in a multivariate trait space rather than on a univariate basis. Such an aim, however, requires considerable theoretical development and may well require substantially more data. For now, however, we should be aware that we are collapsing our trait space, and each of our measured traits is not independent. For example, in our system there is a strong correlation between CTmin and CTmax, thus we should be aware that these two traits should not get equal weighting when we use our traits to rank environmental variables by their importance to local adaptation

#### Neutral clines

Finally, our approach should allow us to identify when geographic variation is a result of genetic variation. That is, it can weed out relationships that are driven by plasticity or habitat choice. Covariation between genotype and environment will often be the result of local adaptation, but can also arise for non-adaptive reasons, the most obvious being trait clines caused by the historical spread of population (62). In principle, and again, with careful attention to sample design (i.e., a sample design which minimizes the covariation between space and environment), it should be possible to separate spatial from environmental patterns.

### Conclusion

There is increasing urgency to identify populations that will act as suitable sources for targeted gene flow efforts in the face of climate change. To identify these populations, we need to know which traits influence sensitivity to climate and are locally adapted. Traditional approaches to unearthing local adaptation (reciprocal transplants and common garden experiments) are time consuming, and often cannot attribute adaptation to any particular environmental driver. Local adaptation is, however, undermined by gene flow, and we should be able to use this fact to sort patterns of local adaptation from patterns with other causes. Here we have demonstrated this approach: using connectivity as a proxy for gene flow, and looking for its effect on trait–environment relationships. Our analysis, using a species of lizard from Australia's Wet Tropics rainforest, suggests the approach has merit: the results we achieve are coherent and suggest local adaptation is the overwhelming signal in the set of trait–environment relationships tested. As well as implying a strong role for local adaptation, we have effectively ranked environmental drivers of local adaptation, finding evidence that precipitation and temperature are important environmental variables with regard to local adaptation in our system. Our analysis also suggests that some traits exhibit strong plastic responses to the environment, particularly in response to precipitation of the driest quarter and the seasonality of temperature and precipitation. These specific results will likely apply to other species that are phylogenetically or ecologically similar to our focal species, but the method has the potential to apply much more broadly. Analytical and sampling challenges remain, however, and we point to avenues whereby the method can be improved. Given the potential of this method to provide evidence of local adaptation, and to provide rapid ranking of the climatic drivers of local adaptation, assessment of the method in a broader array of systems is warranted.

## Acknowledgments

This work was funded by the Tropical Landscapes Joint Venture, the Wet Tropics Management Authority, an Australian Postgraduate Award to SLM, and the Australian Research Council (DP130100318, FT160100198). All physiological data used in these analyses were collected by Amberlee Hatcher at James Cook University. The authors are extremely grateful for her patience and attention to detail while undertaking this task.

## References

1. Reznick DN & Ghalambor CK (2001) The population ecology of contemporary adaptations: what empirical studies reveal about the conditions that promote adaptive evolution. Genetica 112:183–198.

2. Reznick DN, Shaw FH, Rodd FH, & Shaw RG (1997) Evaluation of the rate of evolution in natural populations of guppies (*Poecilia reticulata*). Science 275(5308):1934–1937.

3. Losos JB, Warheit KI, & Schoener TW (1997) Adaptive differentiation following experimental island colonization in *Anolis* lizards. Nature 387(6628):70–73.

4. Stuart YE, et al. (2014) Rapid evolution of a native species following invasion by a congener. Science 346(6208):463–466.

5. Richardson JL, Urban MC, Bolnick DI, & Skelly DK (2014) Microgeographic adaptation and the spatial scale of evolution. Trends Ecol Evol 29(3):165–176.

6. Stockwell CA, Hendry AP, & Kinnison MT (2003) Contemporary evolution meets conservation biology. Trends Ecol. Evol. 18(2):94–101.

7. Kelly MW, Sanford E, & Grosberg RK (2012) Limited potential for adaptation to climate change in a broadly distributed marine crustacean. Proceedings of the Royal Society B-Biological Sciences 279(1727):349–356.

8. Olsson M & Uller T (2003) Thermal environment, survival and local adaptation in the common frog, *Rana temporaria*. Evol Ecol Res 5(3):431–437.

9. Sinervo B, et al. (2010) Erosion of Lizard Diversity by Climate Change and Altered Thermal Niches. Science 328(5980):894–899.

10. Hoffmann AA & Sgrò CM (2011) Climate change and evolutionary adaptation. Nature 470(7335):479–485.

11. Kawecki TJ & Ebert D (2004) Conceptual issues in local adaptation. Ecol Lett 7(12):1225–1241.

12. Reznick DN & Travis J (1996) The empirical study of adaptation in natural populations. Adaptation, (Academic Press, San Diego, CA), pp 243–289.

13. Blanquart F, Kaltz O, Nuismer SL, & Gandon S (2013) A practical guide to measuring local adaptation. Ecology Letters 16:1195–1205.

14. Wright JW, Stanton ML, & Scherson R (2006) Local adaptation to serpentine and non-serpentine soils in *Collinsia sparsiflora*. Evol Ecol Res 8(1):1–21.

15. García-Ramos G & Kirkpatrick M (1997) Genetic models of adaptation and gene flow in peripheral populations. Evolution 51(1):21–28.

16. Wright S (1931) Evolution in Mendelian populations. Genetics 16(2):0097–0159.

17. Dhondt AA, Adriaensen F, Matthysen E, & Kempenaers B (1990) Nonadaptive Clutch Sizes in Tits. Nature 348(6303):723–725.

18. Storfer A, Cross J, Rush V, & Caruso J (1999) Adaptive coloration and gene flow as a constraint to local adaptation in the streamside salamander, *Ambystoma barbouri*. Evolution 53(3):889–898.

19. Sandoval CP (1994) The Effects of the Relative Geographic Scales of Gene Flow and Selection on Morph Frequencies in the Walking-Stick *Timema christinae*. Evolution 48(6):1866–1879.

20. IPCC (2014) Climate Change 2014: Synthesis Report. Contribution of Working Groups I, II and III to the Fifth Assessment Report of the Intergovernmental Panel on Climate Change (IPCC, Geneva, Switzerland) p 151.

21. Meehl GA, et al. (2007) Global Climate Projections. in Climate Change 2007: The Physical Science Basis. Contribution of Working Group I to the Fourth Assessment Report of the Intergovernmental Panel on Climate Change, eds Solomon S, Qin D Manning M Chen Z Marquis M Averyt KB, Tignor M & Miller HL (Cambridge University Press, Cambridge, United Kingdom and New York, NY, USA).

22. Aitken SN & Whitlock MC (2013) Assisted gene flow to facilitate local adaptation to climate change. Annu Rev Ecol Evol Syst 44:367–388.

23. Williams SE, Shoo LP, Isaac JL, Hoffmann AA, & Langham G (2008) Towards an integrated framework for assessing the vulnerability of species to climate change. PLoS Biol 6(12):2621–2626.

24. Weeks AR, et al. (2011) Assessing the benefits and risks of translocations in changing environments: a genetic perspective. Evolutionary Applications 4(6):709–725.

25. Kelly EL & Phillips BL (2016) Targeted gene flow for conservation. Cons. Biol. 30:259–267.

26. Wilson SK & Swan G (2010) A complete guide to reptiles of Australia (Reed New Holland, Sydney, N.S.W.) p 512 pp.

27. Llewelyn J, Macdonald S, Hatcher A, Moritz C, & Phillips BL (2016) Intraspecific variation in climate-relevant traits in a tropical rainforest skink. Diversity and Distributions 22(10):1000–1012.

28. Phillips BL, Llewelyn J, Hatcher A, Macdonald S, & Moritz C (2014) Do evolutionary constraints on thermal performance manifest at different organizational scales? J. Evol. Biol. 27:2687–2694.

29. Easterling DR, et al. (2000) Climate extremes: Observations, modeling, and impacts. Science 289(5487):2068–2074.

30. Parmesan C, Root TL, & Willig MR (2000) Impacts of extreme weather and climate on terrestrial biota. B Am Meteorol Soc 81(3):443–450.

31. Macdonald SL, Llewelyn J, Moritz C, & Phillips BL (2017) Peripheral isolates as sources of adaptive diversity under climate change. Frontiers in Ecology and Evolution 5.

32. R Core Team (2015) R: A Language and Environment for Statistical Computing (R Foundation for Statistical Computing, Vienna, Austria).

33. Llewelyn J, Macdonald SL, Hatcher A, Moritz C, & Phillips BL (submitted) Intraspecific variation in climate-relevant traits in a tropical rainforest skink.

34. Wegener JE, Gartner GEA, & Losos JB (2014) Lizard scales in an adaptive radiation: variation in scale number follows climatic and structural habitat diversity in *Anolis* lizards. Biol J Linn Soc 113(2):570–579.

35. Klaczko J, Ingram T, & Losos J (2015) Genitals evolve faster than other traits in *Anolis* lizards. Journal of Zoology 295(1):44–48.

36. Bogert CM (1949) Thermoregulation in Reptiles, a Factor in Evolution. Evolution 3(3):195–211.

37. Hoffmann AA (2010) Physiological climatic limits in *Drosophila*: patterns and implications. J Exp Biol 213(6):870–880.

38. Clusella-Trullas S, Blackburn TM, & Chown SL (2011) Climatic Predictors of Temperature Performance Curve Parameters in Ectotherms Imply Complex Responses to Climate Change. Am Nat 177(6):738–751.

39. Angilletta MJ (2009) Thermal adaptation: a theoretical and empirical synthesis (Oxford University Press, Oxford; New York) pp xii, 289 p. 281 col. plate.

40. Goodman BA, Miles DB, & Schwarzkopf L (2008) Life on the Rocks: Habitat Use Drives Morphological and Performance Evolution in Lizards. Ecology 89(12):3462–3471.

41. Huey RB & Webster TP (1976) Thermal Biology of *Anolis* Lizards in a Complex Fauna: the *cristatellus* Group on Puerto-Rico. Ecology 57(5):985–994.

42. Crowley SR (1987) The Effect of Desiccation Upon the Preferred Body-Temperature and Activity Level of the Lizard *Sceloporus indulatus*. Copeia (1):25–32.

43. Jones SM, Waldschmidt SR, & Potvin MA (1987) An Experimental Manipulation of Food and Water - Growth and Time-Space Utilization of Hatchling Lizards (*Sceloporus undulatus*). Oecologia 73(1):53–59.

44. Lorenzon P, Clobert J, Oppliger A, & John-Alder H (1999) Effect of water constraint on growth rate, activity and body temperature of yearling common lizard (*Lacerta vivipara*). Oecologia 118(4):423–430.

45. Stamps J & Tanaka S (1981) The Influence of Food and Water on Growth-Rates in a Tropical Lizard (*Anolis aeneus*). Ecology 62(1):33–40.

46. Myers VI & Heilman MD (1969) Thermal Infrared for Soil Temperature Studies. Photogramm Eng 35(10):1024–1032.

47. Martin TL & Huey RB (2008) Why “Suboptimal” is optimal: Jensen’s inequality and ectotherm thermal preferences. Am Nat 171(3):E102–E118.

48. Bonebrake TC & Mastrandrea MD (2010) Tolerance adaptation and precipitation changes complicate latitudinal patterns of climate change impacts. Proc Natl Acad Sci U S A 107(28):12581–12586.

49. Kingsolver JG & Huey RB (1998) Evolutionary analyses of morphological and physiological plasticity in thermally variable environments. Am Zool 38(3):545–560.

50. Gilchrist GW (1995) Specialists and Generalists in Changing Environments. I. Fitness Landscapes of Thermal Sensitivity. Am Nat 146(2):252–270.

51. Phillips BL, et al. (2016) Heat hardening in a tropical lizard: geographic variation explained by the predictability and variance in environmental temperatures. Func. Ecol. 30:1161–1168.

52. Polechová J, Barton N, & Marion G (2009) Species' range: adaptation in space and time. Am Nat 174(5):E186–E204.

53. Seebacher F (2005) A review of thermoregulation and physiological performance in reptiles: what is the role of phenotypic flexibility? Journal of comparative physiology. B, Biochemical, systemic, and environmental physiology 175(7):453–461.

54. Llewelyn J, Macdonald S, Moritz C, & Phillips BL (2017) Thermoregulatory behaviour explains countergradient variation in the upper thermal limit of a rainforest skink. Oikos 126(5):748–757.

55. DERM (2011) Queensland Department of Environment and Resource Management, regional ecosystem mapping version 7.0.

56. Singhal S & Moritz C (2012) Strong selection against hybrids maintains a narrow contact zone between morphologically cryptic lineages in a rainforest lizard. Evolution 66(5):1474–1489.

57. Palmer SCF, Coulon A, & Travis JMJ (2011) Introducing a 'stochastic movement simulator’ for estimating habitat connectivity. Methods in Ecology and Evolution 2(3):258–268.

58. Petkova D, Novembre J, & Stephens M (2016) Visualizing spatial population structure with estimated effective migration surfaces. Nature Genetics 48(1):94–100.

59. Merila J, Sheldon BC, & Kruuk LEB (2001) Explaining stasis: microevolutionary studies in natural populations. Genetica 112:199–222.

60. Lynch M & Walsh B (1998) Genetics and Analysis of Quantitative Traits (Sinauer Associates, Sunderland, MA).

61. Blows MW (2007) A tale of two matrices: multivariate approaches in evolutionary biology. J. Evol. Biol. 20(1):1–8.

62. Phillips BL, Brown GP, & Shine R (2010) Life-history evolution in range-shifting populations. Ecology 91(6):1617–1627.

63. Jones, D.A., Wang, W. & Fawcett, R. (2009) High-quality spatial climate data-sets for Australia. Australian Meteorological and Oceanographic Journal, 58, 233–248.

64. McMahon, J.P., Hutchinson, M.F., Nix, H.A. & Ord, K.D. (1995). yANUCLIM user's guide. Centre for Resource and environmental Studies, Australian National University, Canberra.

65. Storlie, C.J., Phillips, B.L., VanDerWal, J.J. & Williams, S.E. (2013) Improved spatial estimates of climate predict patchier species distributions. Diversity and Distributions, 19, 1106–1113.

66. Graham, Catherine H., VanDerWal, Jeremy, Phillips, Steven J., Moritz, Craig, and Williams, Stephen E. (2010). Dynamic refugia and species persistence: tracking spatial shifts in habitat through time. Ecography, 33(6):1062–1069.

